# GlycoRNA-L and glycoRNA-S mediate human monocyte adhesion via binding to Siglec-5

**DOI:** 10.1101/2024.08.27.609838

**Authors:** Mingui Fu, Yisong Qian, Evan Huang, Zain Schwarz, Hannah Tai, Katherine Tillock, Tianhua Lei

## Abstract

It was recently reported that RNAs can be glycosylated, and a majority of such glycosylated RNAs (referred to as glycoRNAs) are located on the outer cell surface. We here reported that there are two forms of glycoRNAs, named as glycoRNA-L and glycoRNA-S, robustly expressed in human monocytes. Both of glycoRNA-L and glycoRNA-S contributed to the interaction of human monocytes and endothelial cells via directly binding to Siglec-5. GlycoRNA-L predominantly expressed in most of tissues and cell lines. GlycoRNA-S only expressed in some cell lines and tissues. Siglec-5 preferentially binds to glycoRNA-L than glycoRNA-S. The composition of glycan chains in glycoRNA-L and glycoRNA-S is different. GlycoRNA-L contains more sialic acid, whereas glycoRNA-S contains more GlcNAc. Together, these results demonstrate that two forms of glycoRNAs exist, which may play significant role in controlling the interaction of human monocytes and endothelial cells and contribute to the pathogenesis of inflammatory diseases.

## INTRODUCTION

There are many biomolecules localized on outer cell surface including proteins, lipids and glycans, which are important to mediate cell-cell interaction and play important roles in inflammatory processes. Typically, RNAs are not expected to be present on the surface of cells. Several previous studies have observed the existence of stable RNA-cell surface interactions (1-3). The first membrane-bound RNA was discovered in bacteria (3), and it is a noncoding RNA transcribed from the same operon encoding a transmembrane protein. Following co-transcription, the noncoding RNA and the transmembrane protein assemble into a ribonucleoprotein that gets incorporated into the cell membrane (3). Intriguingly, several recent studies identified varieties of RNA molecules localized on cell plasma membrane. Huang et al. identified a group of RNA molecules that are stably attached to outer cell surface. All the RNA molecules they identified are RNA fragments belong to long noncoding RNAs (lncRNAs) or mRNAs (4). Flynn et al. have reported that RNAs can be glycosylated, and a majority of such glycosylated RNA (called glycoRNAs) are located on the outer cell surfaces including human stem cells and several cancer cell lines (5). Zhang et al. further demonstrated that the cell surface glycoRNAs are important molecules that mediate neutrophils recruitment into inflammatory area by selective interaction with adhesion molecules on vascular endothelial cells such as P-selectin (6). Although there are still many things unclear about the newly discovered cell surface RNAs, evidence suggests that there are small noncoding RNAs (snRNAs), which are synthesized inside cells and transported to cell plasma membrane. These intriguing discoveries open a new avenue for studying the molecular mechanisms that control inflammation and understanding the molecular controls of the pathogenesis of inflammatory diseases.

The pathogenesis of inflammatory diseases is not completely clear but is exacerbated by excessive infiltration of inflammatory cells including neutrophils and monocytes (7, 8). The inflammatory cells respond quickly to infection or tissue injuries by migrating from the circulation toward inflammatory sites, a process involving many cell-cell interactions. Lectin-like adhesion glycoproteins, called selectins, mediate leukocyte rolling, while the firm adhesion and subsequent transendothelial migration of leukocytes are mediated by the interaction of integrins on leukocytes with immunoglobin-like adhesion molecules on endothelial cells (ICAM-1 and VCAM-1, 9-11). However, the molecular mechanisms controlling inflammatory cells adhesion and infiltration remain incompletely understood. Targeting the process of inflammatory cell adhesion and infiltration to treat inflammatory diseases is still lacking.

Here we reported that there are two forms of glycoRNAs, named as glycoRNA-L and glycoRNA-S, robustly expressed in human monocytes. Both of glycoRNA-L and glycoRNA-S contributed to the adhesion of human monocytes to endothelial cells via directly binding to siglec-5. GlycoRNA-L predominantly expressed in most of tissues and cell lines. However, glycoRNA-S only expressed in some tissues and cell lines including adipose and human monocytes. Siglec-5 preferentially binds to glycoRNA-L than glycoRNA-S. Though the RNA species contained in glycoRNA-L and glycoRNA-S may be similar, the composition of glycan chains in glycoRNA-L and glycoRNA-S is different. Collectively, we found that there are two forms of glycoRNAs expressed in human monocytes, which may play an important role in the regulation of inflammation and become new targets for treatment of inflammatory diseases.

## RESULTS

### Expression of glycoRNA-L and glycoRNA-S in human monocytes

To investigate whether human monocytes express glycoRNAs, we used THP1 cells, a human monocyte cell line, as a model. Following the method described by Flynn et al. (5), we metabolically labeled live THP1 cells using N-azidoacetylmannosamine-tetraacylated (Ac4ManNAz), which can enter the sialic acid synthesis pathway, leading to azide-modified sialic acids in glycans. Purified RNAs from labeled cells were then reacted in vitro with dibenzocyclooctyne-polyethylene-glycol-4-biotin (DBCO-PEG4-biotin) via click chemistry so that sialic-acid-containing glycoRNAs could be biotin-modified. When analyzed by denaturing electrophoresis and subsequent blotting, we observed two bands from the blot (**Figure 1A**). One band was migrated with ∼11 kb RNA, designated as glycoRNA-L; the other band was migrated with ∼0.6 kb RNA, designated as glycoRNA-S. Both glycoRNA-L and glycoRNA-S were abolished upon digestion of purified RNA by RNase prior to electrophoresis. To exclude the glycoRNA-S is an artificial form caused by labeled chemical, we incubated total RNA with 0, 5, 10 and 20 mM of Ac4ManNAz, which is 200 folds than the dose in cells. The RNAs was cleaned up by column and conjugated with biotin followed by northern blot analysis. As shown in **Figure S1A**, there are clearly two bands from in vivo labeled cells, however, in vitro incubation of total RNA with up to 20 mM Ac4ManNAz did not produce any signals. We noted that Flynn et al. (5) and Zhang et al. (6) only detected the glycoRNA-L, but not glycoRNA-S. To exclude the possibility that the difference was caused by Ac4ManNAz from different sources, we incubated THP1 cells with Ac4ManNAz from Torcris (we used in this report), Click Chemistry (Flynn et al and Zhang et al, used in their papers) and TargetMol. As shown in **Figure S1B**, though different efficiency, all three Ac4ManNAz can label the glycoRNA-S. We didn’t see the glycoRNA-L in this blot, which may be due to the shorter transfer time. To further confirm that glycoRNA-S is a new form of glycoRNAs, we added glycan synthesis inhibitors with Ac4ManNAz into THP1 cells. As shown in **Figure 1B**, inhibition of oligosaccharyltransferase (OST) by NGI-1 diminished the expression of both glycoRNA-L and glycoRNA-S. However, kifunensine, an inhibitor of the N-glycan trimming enzyme α-mannosidase I, did not efficiently change the expression of glycoRNA-L and glycoRNA-S. Together, the data above support the presence of two forms of glycoRNAs, or more precisely sialoglycoRNAs, in human monocytes.

**Figure 1.**
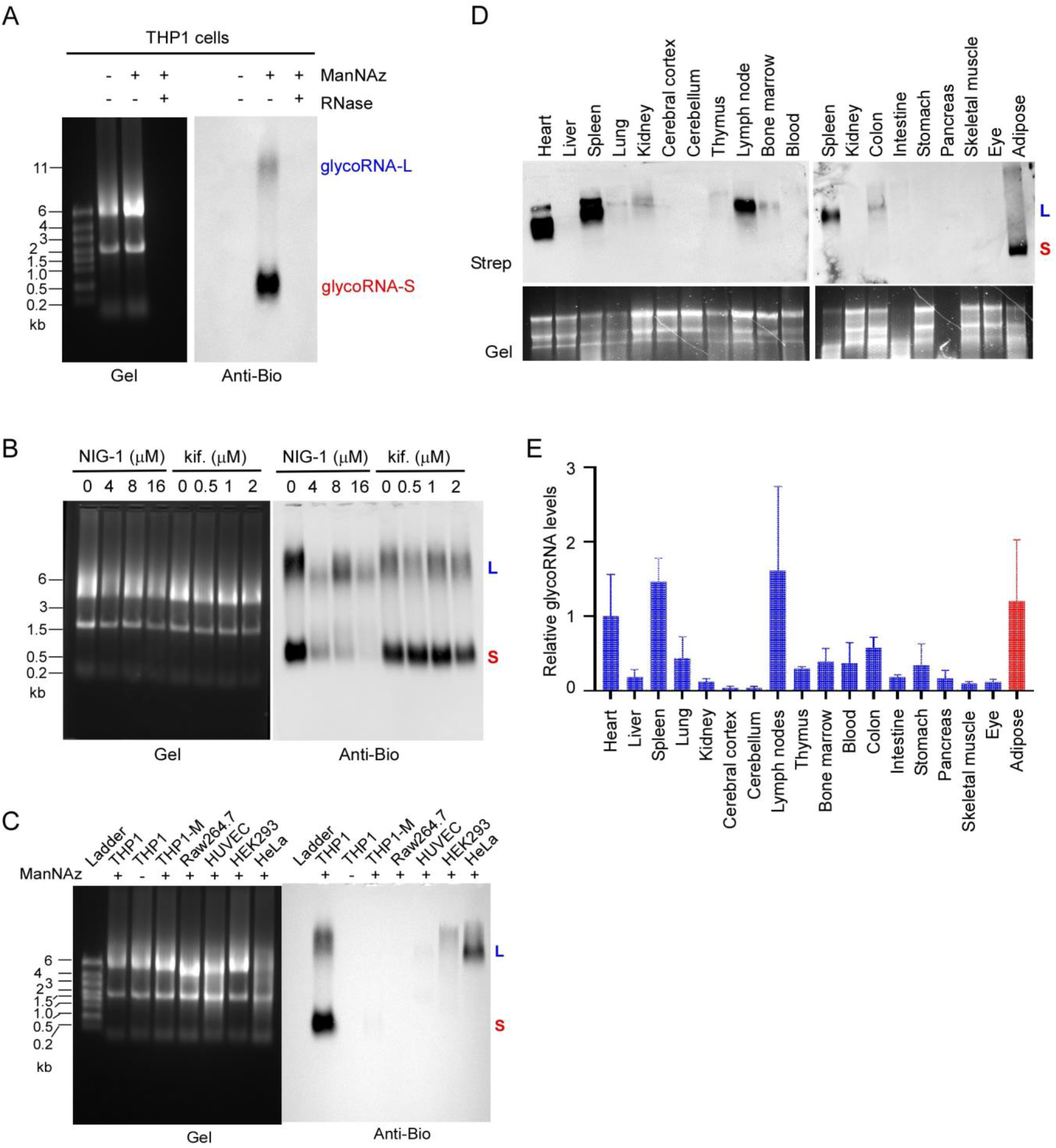
Expression of glycoRNA-L and glycoRNA-S in human monocytes. (A) THP1 cells were treated with or without Ac_4_ManNAz. RNAs were extracted from the cells and treated with or without RNase. Then the RNA samples were reacted with DBCO-PEG4-biotin. RNAs were analyzed on an agarose gel (left) and then blotted with an anti-biotin antibody (right). Representative images are shown. (B) THP1 cells were treated with Ac_4_ManNAz and different doses of NIG-1 or kifunensine as indicated. RNAs were extracted from the cells and analyzed by northern blot. Representative images are shown. (C) RNA from different cells treated with 100 μM Ac_4_ManNAz for 24 hours and then analyzed by the same procedure as above. Representative images are shown. (D) RNA blot of mouse RNA after in vivo Ac_4_ManNAz delivery via intraperitoneal injection for 2 days at 300 mg/kg/day. RNA extracted from the tissues were analyzed by northern blot and probed by streptavidin-HRP. (E) The relative glycoRNA levels from different tissues were analyzed by Image J. Data were presented as mean±sd, n=3. Blue column represents glycoRNA-L, red column represents glycoRNA-S. Also see Figure S1.

To explore the distribution of glycoRNA-L and glycoRNA-S in different cells, we incubated Ac4ManNAz with several human and mouse cell lines as indicated in **Figure 1C** and followed by northern blot analysis as described above. As shown in **Figure 1C**, THP1 cells expressed both glycoRNA-L and glycoRNA-S, whereas HeLa and HEK293 cells only expressed glycoRNA-L. There were no glycoRNA detected in THP1-derived macrophages, Raw264.7, and rest human umbilical vain endothelial cells (HUVECs). Next, we examined the distribution of glycoRNA-L and glycoRNA-S in mouse tissues. Adult C57/BL6 mice were intraperitoneally injected with 300 mg/kg/day of Ac4ManNAz for 2 days. Total RNA was isolated from the tissues as indicated and analyzed by the same procedures as described above. As shown in **Figure 1D**, the glycoRNAs were predominantly expressed in immune organs such as spleen, thymus, bone marrow and lymph nodes. High levels of expression were also observed in heart and adipose. Low levels of expression were observed in lung, colon and kidney. There was no expression of glycoRNAs in brain, eye, muscle, intestine and stomach. Interestedly, though most tissues expressed the glycoRNA-L, adipose expressed glycoRNA-S. Together, the predominant expression of glycoRNAs in immune organs suggests that glycoRNAs may be critical in the regulation of immune response.

### GlycoRNAs mediate the interaction between human monocytes and endothelial cells

As Flynn et al. (5) reported that the majority of glycoRNAs is localized on outer cell surface, next we examined whether RNase incubation with living cells can remove the glycoRNAs from cell surface. Briefly, the labeled THP1 cells as described above were spun down and washed once with Hank-Balanced Buffer Solution (HBSS), then 10^7^ THP1 cells were resuspended in 1 ml HBSS and incubation with 0, 1, 5 μl of RNase cocktail (containing RNase A 0.5 unit/μl and RNase T 20 unit/μl) at 37 °C for 10 min. Then the cells were washed by HBSS and added 1 ml of TRIzol. The total RNAs from the cells were extracted and biotin modified as described above. Technical note that if the total RNA was extracted by the procedure described in **Figure S1C**, the step of proteinase K treatment can be omitted as shown in **Figure S1D**. As shown in **Figure 2A**, the glycoRNAs can be efficiently removed by RNase treatment, which proved that glycoRNAs were expressed on cell surface. The same concentration of RNase treatment did not affect the cell viability, suggesting that RNase did not damage cell membrane and enter cells (**Figure 2B**). To understand if glycoRNAs mediate monocyte adhesion on EC layer, we performed monocyte adhesion assay according to our previous publication (12). Briefly, HUVECs were seeded on 6-well plate and stimulated with or without TNF-α (10 ng/ml) plus LPS (1 μg/ml) for 16 h. THP1 cells were pretreated with 0, 1 or 5 μl of RNase cocktail in 1 ml of cells at 37 °C for 10 mins and then labeled with PHK67 (Sigma) for 5 min at 37°C before being added to HUVECs and co-cultured for 1 h. Non-adherent cells were removed by gently washing with cold RPMI1640 medium. The images of adherent THP1 cells were taken under microscopy. 6 images from different fields were taken from each well and the adherent cell numbers were counted by Qupath in a two-blind manner. First, we verified that activated HUVECs was attached with much more THP1 cells than rest HUVECs (**Figure S2**). As shown in **Figure 2B**, RNase treatment significantly inhibited THP1 cell adherence on activated ECs. These data support that cell surface glycoRNAs are necessary for efficient monocyte-endothelial interactions.

**Figure 2.**
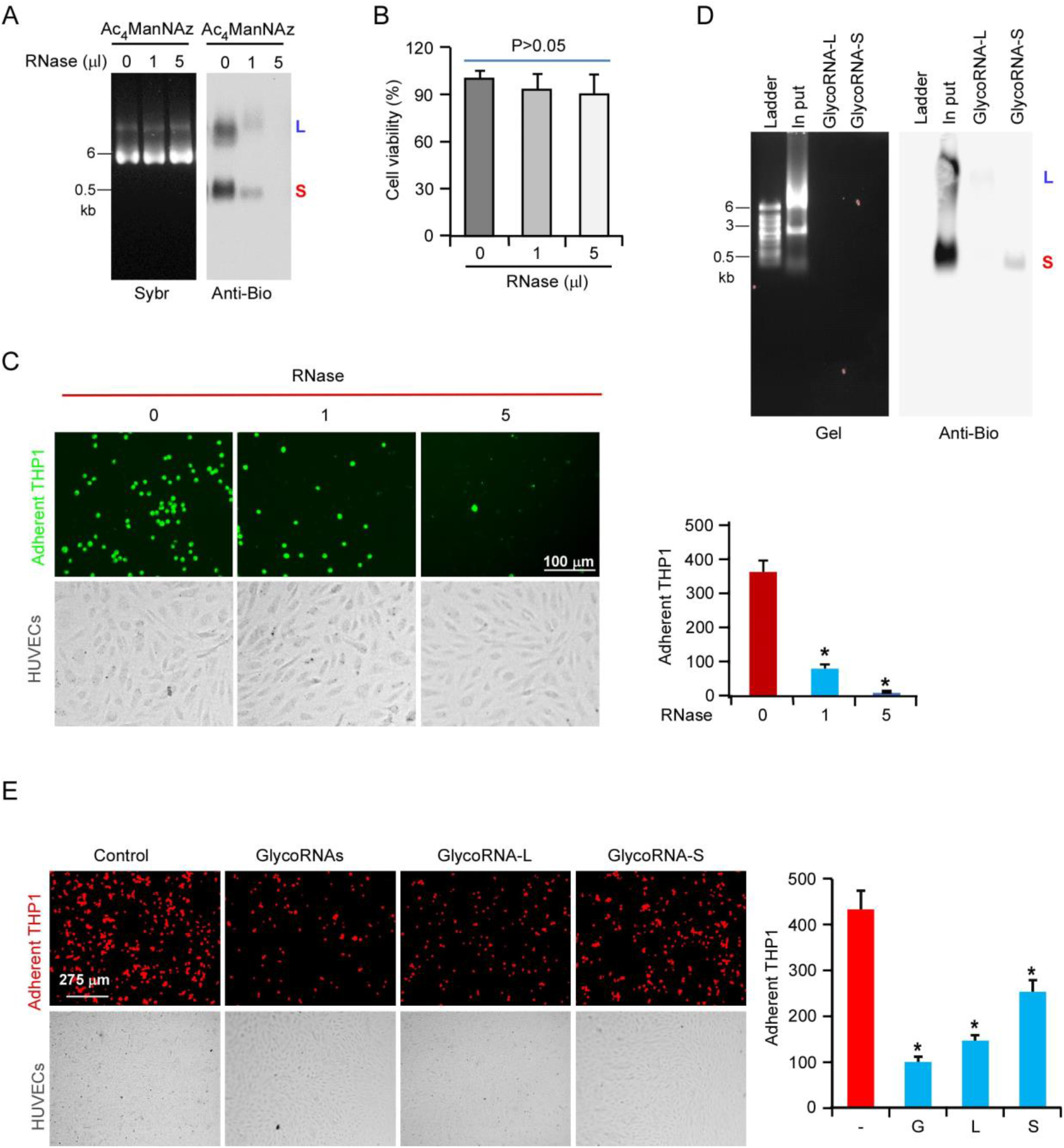
GlycoRNAs mediated human monocytes adhesion on activated endothelial cells. (A) THP1 cells were metabolic labeled by Ac_4_ManNDAz for 24 h. The cells were harvested by centrifuge and washed with HBSS, then resuspended in 1 ml HBSS and incubated with 0, 1, or 5 μl of RNase cocktail. The total RNAs were isolated and analyzed as described above. (B) THP1 cells were subjected to the same treatment with RNase cocktail. The cell viability was analyzed by Trypan blue exclusion test. (C) The cell images taking from RNase-treated and untreated THP1 cells then labeled by PHK67. The labeled THP1 cells incubated with activated HUVECs at 37°C for 1 h, then the adherent THP1 cells were imaged by microscopy. The image represents one fourth of original image to make the cells more visible (left). 6 random fields per well were imaged and the cell numbers in each field were counted by Qupath. The adherent cell numbers were averaged and presented as mean±SD, n=3 (right). (D) Purified glycoRNA-L and glycoRNA-S were analyzed by northern blot as described above. (E) Before adding THP1-RFP cells, the activated HUVECs were pretreated with or without 100 ng/ml of glycoRNAs, glycoRNA-L or glycoRNA-S. Then THP1-RFP cells were added into HUVECs and analyzed by the same procedure described above. See also Figure S2.

To further characterize the role of glycoRNA-L and glycoRNA-S in mediating monocyte adhesion, we purified glycoRNA-L and glycoRNA-S by the procedures described by Flynn et al (5) with some modification. First, the biotinylated glycoRNAs were purified by Myone C1 Streptavidin beads. Then the purified glycoRNAs were separated by low-melting temperature agarose gel and the glycoRNA-L and glycoRNA-S were recovered from the gel slices. As shown in **Figure 2D**, we successfully purified the glycoRNA-L and glycoRNA-S. Then we pretreated the activated HUVECs with glycoRNAs or glycoRNA-L or glycoRNA-S followed by THP1 cell adhesion assay. As shown in **Figure 2E**, both glycoRNA-L and glycoRNA-S contributed to mediating THP1 cell adhesion, but glycoRNA-L may contribute more than glycoRNA-S.

### Identification of Siglec-5 as the binding partner of glycoRNAs from THP1 cells

Next, we searched the target proteins on HUVECs that might bind to the glycoRNAs on THP1 cells. First, we purchased the commercially available endothelial adhesion proteins including P-selectin, E-selectin, VCAM-1, ICAM-1, Siglec-4a and Siglec-5-Fc chimeras. Then we probed their binding to THP1 cells by flow cytometry and microscopy. As shown in **Figure S3A**, among of 6 recombinant proteins tested, only P-selectin, Siglec-4a and Siglec-5 strongly bind to THP1 cells. Then, we pretreated THP1 cells with RNase before incubation with P-selectin, Siglec-4a or Siglec-5-Fc followed by the analysis of flow cytometry and microscopy. As shown in **Figure 3A&B**, only Siglec-5 binding was sensitive to RNase treatment. To further confirm that Siglec-5 directly binds to the glycoRNAs from THP1 cells, we performed immunoprecipitation. Briefly, biotinylated RNAs from ManNAz-labeled THP1 cells were incubated with P-selectin, E-selectin, VCAM-1, Siglec-4a and Siglec-5-Fc chimeras and then pulldown by incubation with anti-Fc agarose beads. The bound biotinylated glycoRNAs were detected by northern blot analysis as described previously. As shown in **Figure 3C**, only Siglec-5 can pulldown glycoRNA-L and glycoRNA-S. Next, we directly probed the northern blot membrane with P-selectin, E-selectin, Siglec-4a and Siglec-5-Fc chimera and anti-human Fc-HRP complex as described by Zhang et al (6). As shown in **Figure 3D** and **Figure S3B**, only Siglec-5 specifically binds to the native glycoRNAs from THP1 cells. Both Siglec-4a and Siglec-5 can bind to the native glycoRNAs from HeLa cells. Interestedly, Siglec-5 preferentially binds to glycoRNA-L in both THP-1 cells and HeLa cells. Both P-selectin and E-selectin did not bind to the glycoRNAs from THP1, HL60, HUVECs, HEK293 and HeLa cells (**Figure S3B**). It appeared that P-selectin non-specially bind to 28S RNA.

**Figure 3.**
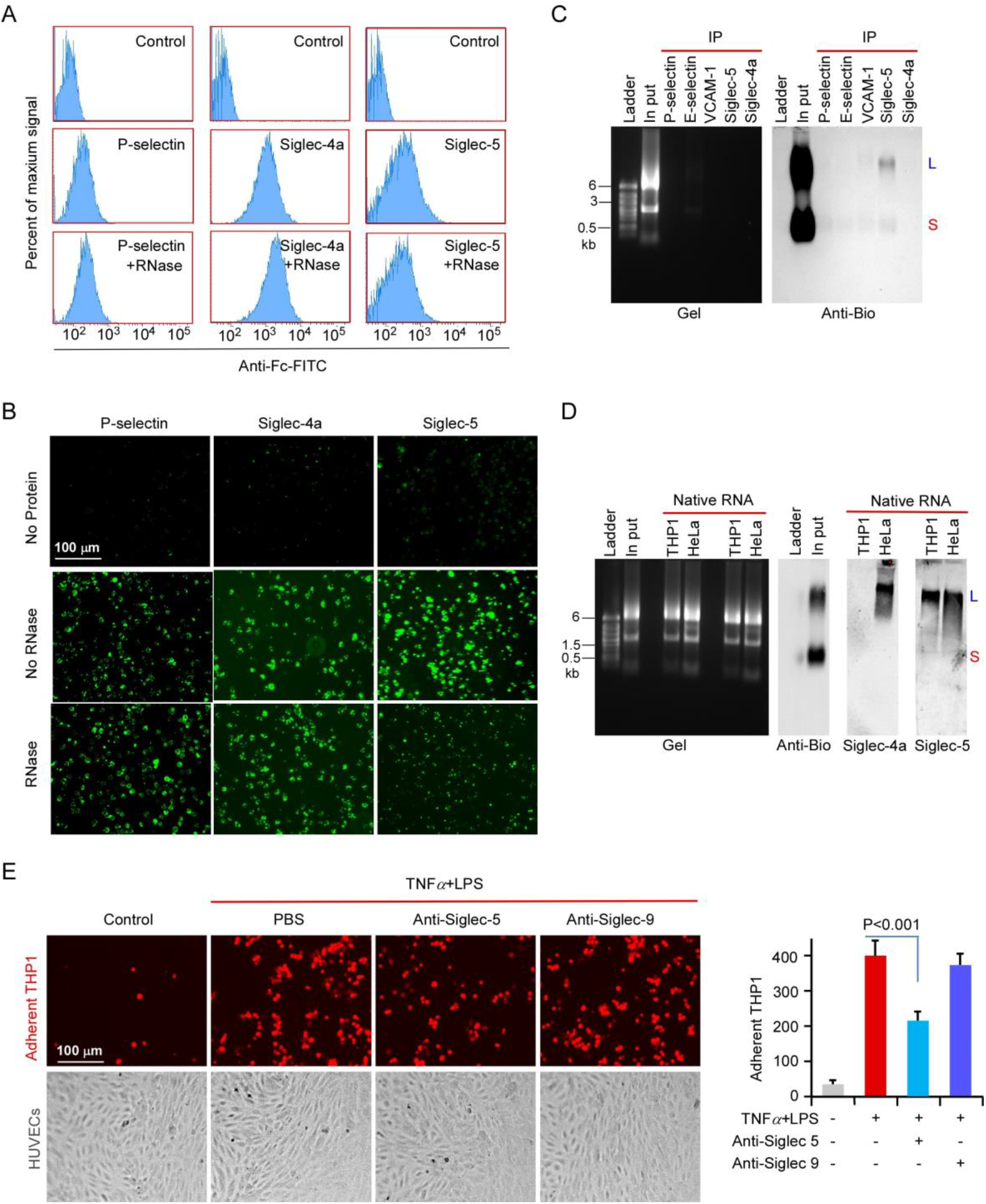
Identification of Siglec-5 as the binding partner of glycoRNAs from THP1 cells. (A) FACS analysis of THP1 cells pretreated with or without RNase then stained with the indicated pre-complex. (B) The images taken from these cells in (A) by microscopy. (C) Biotinylated RNA was incubated with pre-binding anti-Fc-beads with indicated protein-Fc chimera and then analyzed by northern blot as described above. (D) Native RNAs were analyzed by northern blot. After blocking, the membrane was probed by pre-complex of Siglec-4a-Fc (1.5 μg/ml) or Siglec-5-Fc (1.5 μg/ml) with anti-Fc-HRP (1.5 μg/ml) for 2 h at 4°C. The blot was imaged by iBright 1500. Blot of biotinylated RNA from labeled THP1 cells was probed by anti-biotin-HRP. (E) Before adding THP1-RFP cells, the activated HUVECs were pretreated with or without 1.5 μg/ml of anti-siglec-4a or anti-Siglec-5. Then THP1-RFP cell adherent was analyzed by the same procedure described above. See also Figure S3.

Next, we tested whether Siglec-5 antibody can block the adhesion of THP1 cells on activated HUVECs. Before incubation with THP1 cells, the activated HUVECs were pretreated with anti-Siglec-5 or anti-Siglec-9. As shown in **Figure 3E**, treatment with anti-Siglec-5 but not anti-Siglec-9 significantly attenuated the THP1 cell adhesion. These results suggest that the binding partner of Siglec-5 and glycoRNAs mediates the interaction of THP1 cells and HUVECs. To examine the expression of Siglec-5 on HUVECs, HUVECs were treated with TNFα and LPS for different times as indicated in **Figure S3C**. The cell lysates were subjected to western blot analysis with the antibodies as indicated. As shown in **Figure S3C**, VCAM-1 was significantly induced by TNFα and LPS treatment as expected. However, Siglec-5 was constitutively expressed in rest HUVECs and slightly increased (∼2 folds) after stimulation with TNFα and LPS.

### Characterization of the structure of glycoRNA-L and glycoRNA-S

Next, we try to characterize the structure of glycoRNA-L and glycoRNA-S. As previous report suggested that glycoRNAs are synthesized in vivo by conjugating the glycan chains with the fragments from non-coding small RNAs (5). A recent report demonstrated that glycan chain is attached to the RNA through covalently binding to 3-(3-amino-3-carboxypropyl)uridine (acp^3^U, ref. 13). In mammalian, the glycan chains on glycoproteins and glycolipids are commonly composed by sialic acid, galactose, glucose, GalNAc, GlcNAc, mannose, fructose and xylose (14). To determine the composition of glycan chains on glycoRNA-L and glycoRNA-S, we purchased four commercially available azido-sugar molecules. We labeled the THP1 and HeLa cells with these probes and analyzed by northern blot as described above. As shown in **Figure 4A**, glycoRNA-L is heavily labeled by ManNAz, whereas glycoRNA-S was labeled by ManNAz and GlcNAz. These results suggest that the composition of glycan chains on glycoRNA-L and glycoRNA-S is different. At least, glycoRNA-L has more sialic acid than glycoRNA-S, whereas glycoRNA-S has more GlcNAc than glycoRNA-L. Interestedly, we observed a new form of glycoRNA, named as glycoRNA-M, as it migrates as ∼1.0 kb and at the middle between glycoRNA-L and glycoRNA-S. This glycoRNA is heavily labeled by GalNAz, suggesting that it may be conjugated by more than one GalNAc molecules. We observed a weak band in FucAz-labeled cells. The size is around 5.0 kb, which is similar to the band resulted by chemically catalyzed in vitro in Flynn’s paper. In HeLa cells, only glycoRNA-L was labeled by ManNAz, GalNAz and GlcNAz. There is no glycoRNA-S. However, glycoRNA-M is heavily labeled by GalNAz like that in THP1 cells (**Figure 4B**). Taken together, these results suggest that the glycan composition between glycoRNA-L and glycoRNA-S is different, which may make their function also different. In addition, in different cells, the glycan composition on glycoRNAs is also different, which may make them have specific function. The discovery of glycoRNA-M suggests that there is a family of glycoRNAs exist based on the distinguish of their glycan chains. Further study is necessary to identify the whole family of glycoRNAs in mammalian cells. Zhang et al reported that after PNGase F digest, the RNA species in glycoRNAs were migrated as a single band around 40 bp (6). We reason that the RNA portions on glycoRNA-L and glycoRNA-S may be similar. As shown in **Figure 4C**, removing the RNA portion by RNase only slightly affect the migration of both glycoRNA-L and glycoRNA-S, which suggest that the migration difference between glycoRNA-L and glycoRNA-S is majorly determined by their glycan chains but not RNA fragments.

**Figure 4.**
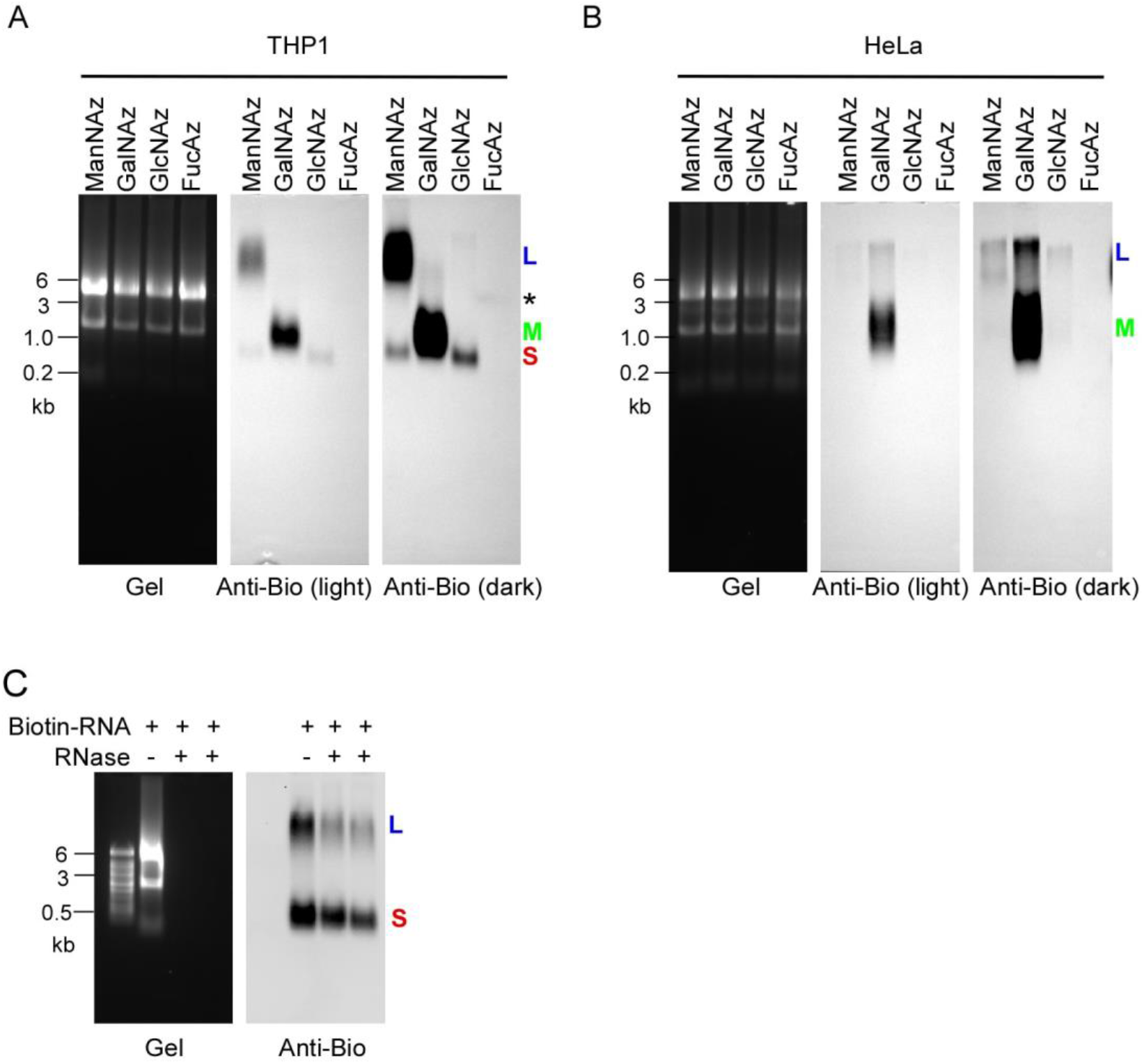
Identification of Siglec-5 as the binding partner of glycoRNAs from THP1 cells. (A) THP1 cells were labeled by 100 μM of Ac_4_ManNAz, GalNAz, GlcNAz and FucAz for 24 hours respectively. RNAs were extracted from the cells. Then the RNA samples were reacted with DBCO-PEG4-biotin. RNAs were analyzed on an agarose gel (left) and then blotted with an anti-biotin antibody (right). Representative images are shown. (B) Blot of RNA from HeLa cells labeled by same reagents as above. (C) 10 μg biotinylated RNA from ManNAz-labeled THP1 cells were treated with 1 μl RNase cocktail and then directly loaded on the denaturing gel and analyzed by same procedure as above.

## DISCUSSION

In this study, we demonstrate that there are two forms of glycoRNAs, or more precisely sialoglyRNAs, robustly expressed in human monocytes. The glycoRNA-L was migrated with RNA around 11 kb and glycoRNA-S was migrated with RNA around 0.6 kb. Actually, these two forms of glycoRNAs also appeared on the blots in several previous reports (5, 15, 16). As the small band is much weak in their blots, the band was thought as a non-specific band catalyzed by labelling chemicals. In my experiments, incubation of total RNAs with 200 folds of concentration of labelling ManNAz did not generate the small band. We also noted that the band generated by ManNAz in vitro in Flynn’s report (5) is around 5 kb, which size is different with glycoRNA-S. Finally, treatment of cells with OST inhibitor abolished both glycoRNA-L and glycoRNA-S, which further demonstrated that glycoRNA-S is a new form of glycoRNAs, but not a non-specific conjugate resulted from labelling chemicals.

The expression of glycoRNA-L is broader than glycoRNA-S. After reviewing all of reports published so far (5, 6, 15-18), there are 16 different cell lines expressed glycoRNA-L including H9, 4188, HeLa, CHO, HOXB8, mouse bone marrow-derived neutrophils, THP1, HEK293, U2OS, human primary alveolar epithelial cells (hPAEpC), HUH7, HAP1, human primary T cells, B cells, monocytes and neutrophils. However, there are only 4 cell lines expressed glycoRNA-S including THP1, 4188, GM78, human primary monocytes. Interestingly, we found that glycoRNAs are abundantly expressed in immune organs in mice such as thymus, lymph nodes, spleen and bone marrow, which suggest that glycoRNAs may significantly involve in the regulation of immune cell functions. We also observed that most tissues only expressed glycoRNA-L and only adipose expressed glycoRNA-S.

The interaction of leukocytes with endothelial cells is the first step of inflammatory response during infection or tissue injury. Glycan-conjugated molecules usually mediate leukocyte adhesion by recognizing their receptors on endothelial cells (19). Our results demonstrate that both glycoRNA-L and glycoRNA-S may involve in mediating the interaction of human monocytes and endothelial cells. We identified Siglec-5 as the receptor for glycoRNA-L and glycoRNA-S. Siglec-5 is a CD33-related Siglecs. It is highly expressed in neutrophils, monocytes and B cells (20). It prefers to bind to sialic acid-containing glycan-conjugated molecules (21). As glycoRNA-L is heavily labeled with sialic acid, it is not surprised that Siglec-5 strongly binds to glycoRNA-L than glycoRNA-S. In our blocking experiment, glycoRNA-L is stronger to inhibit THP1 cell adhesion than glycoRNA-S. Our western blot showed that Siglec-5 was constitutively expressed in HUVECs and its expression was only slightly increased by ∼2 folds. These results cannot explain why THP1 cells did not bind to unstimulated endothelial cells. We hypothesize that there must be a co-factor to assist the interaction of glycoRNAs and Siglec-5 and the expression of the co-factor would be dependent on the activation of endothelial cells.

As Zhang et al. suggest that the glycan portion but not RNA portion mediates cell interaction (6), resolving the composition and structure of glycan chains of glycoRNAs is very important to understand their function. Our results suggest that the sugar composition in the glycan chains of glycoRNA-L is different with that of glycoRNA-S. It is postulated that the different migration nature of glycoRNA-L and glycoRNA-S is due to the glycan chains but not the RNA portion. Indeed, after removing the RNA portion, the migration of glycoRNA-L and glycoRNA-S is a little faster. We hypothesize that RNA portion acts as a “arm” and glycan portion acts as a “hand”. The glycan portions are various as they need to “catch” different cells. However, does the RNA portion also matter? Are the RNA species also important for their function, which need to be further investigated.

From a broader picture, there are many RNA molecules localized on cell surface. Some RNA molecules are modified by glycans, the many others may be modified by other molecules. For example, Huang et al. identified a group of RNA molecules on cell surface, which is mainly composed by RNA fragments from long non-coding RNAs and mRNAs (4). At least two groups generated RNA libraries from lectin-purified glycoRNAs did not contain those RNA species (5, 6), suggesting that those RNA molecules may be modified by other conjugates but not glycans. If focusing on glycoRNA family, there should be more forms of glycoRNAs exist, not only sialic acid-containing glycoRNAs. For example, our results suggest that beside glycoRNA-L and glycoRNA-S, another form of glycoRNA (named as glycoRNA-M) is heavily labeled with GalNAz, but not ManNAz in THP1 and HeLa cells. Identification of the whole glycoRNA family would be important to understand their physiological function and pathological involvement.

In summary, here we report identification of two glycoRNA forms in human monocytes. By serial experiments, we investigated their expression, function, mechanisms and sugar compositions. The limitation of these studies is obvious. First, in vivo functional significance of the two forms of glycoRNAs is not addressed. Second, the structure of glycoRNA-L and glycoRNA-S was not completely elucidated. More studies are needed to further characterize their functions and structures.

## METHODS

### Cell culture

All cells were grown at 37°C and 5% CO_2_. HeLa, HEK293T and Raw264.7 cells were cultured in DMEM media supplemented with 10% fetal bovine serum (FBS) and 1% penicillin/streptomycin (P/S). THP1 and THP1-RFP cells were cultured in RPMI-1640 media with 1 mM HEPES, 1 mM Sodium Pyruvate, 0.001% β-ME and glutamine supplemented with 10% FBS. HUVECs were cultured in EBM2 growth media (Lonza). THP1-derived macrophages were from THP1 cells induced by PMA 200 ng/ml for 5 days and cultured in the same medium with THP1 cells.

### Metabolic labeling of the cells

Stocks of N-azidoacetylmannosamine-tetraacylated (Ac_4_ManNAz, Tocris Bioscience) were made to 500 mM in sterile dimethyl sulfoxide (DMSO). In cell experiments ManNAz was used at a final concentration of 100 μM. *In vitro* experiments with ManNAz used 0, 5, 10, or 20 mM ManNAz (up to 200x the in-cell concentrations) for 2 h at 37°C. Working stocks of glycan-biosynthesis inhibitors were all made in DMSO at the following concentrations and stored at −80°C: 10 mM NGI-1 (Sigma), 10 mM Kifunensine (Kif, Sigma), All compounds were used on cells for 24 h and added simultaneously with ManNAz for labeling. GalNAz, GlcNAz and FucAz were all made to 500 mM stock and used at a final concentration of 100 μM.

### Metabolic labeling in mouse models

All experiments were performed according to guidelines established by the Nanchang University IACUC committee. C57BL/6 mice were crossed and bred in house. ManNAz was prepared by dissolving 100 mg ManNAz in 830 μL 70% DMSO in phosphate buffered saline (PBS), warming to 37°C for 5 min, and then sterile filtering using 0.22 μm Ultra free MC Centrifugal Filter units (Fisher Scientific); this solution was stored at −20°C. Male C57BL/6 mice (8-12 weeks old) were injected once-daily, intraperitoneally with 100 μL of ManNAz (dosed to 300 mg ManNAz/kg/day), while control mice received the vehicle alone. After 2 days, mice were euthanized, and their tissues were harvested. The organs were pressed through a nylon cell strainer and resuspended with PBS to create a single cell suspension. RNA was collected as described below.

### RNA extraction and purification strategies

For total RNA isolation, TRIzol reagent (Thermo Fisher Scientific) was always used as a first step to lyse and denature cells or tissues. After homogenization in TRIzol by pipetting, samples were incubated at 37°C for 10 mins to further denature non-covalent interactions. Phase separation was initiated by adding 0.2x volumes of 100% chloroform, vortexing to mix well, and finally spinning down at 12,000x g for 15 min at 4°C. The aqueous phase was carefully removed, transferred to a fresh tube and mixed with equal volume of isopropanol. The mixture was put in 4°C for 10 mins and then centrifuged at 12,000x g for 10 min at 4°C. The RNA pallet was washed by 1 ml of 75% ethanol (EtOH). The RNA pellet was dry at room temperature and dissolved by RNase-free water. As shown in Figure S1D, if the total RNA was isolated by the procedure described above (**Figure S1C**), the step for proteinase K treatment can be omitted. After enzymatic treatment or biotin-conjugating, the samples were always purified by Zymo column.

### Enzymatic treatment of RNA samples and cells

To digest RNA the following was used: 1 μL of RNase cocktail (0.5U/μL RNaseA and 20U/μL RNase T1, Thermo Fisher Scientific) was added into 20 μl of total RNA at 37 °C for 10 mins. The reaction mixture was purified by Zymo column (**Figure 1A**). To digest biotinylated RNA, 1 μL of RNase cocktail was added into 20 μl of biotinylated RNA 37 °C for 10 mins. The reaction mixture was directly mixed with equal volume of dfGLB II for northern blot analysis (**Figure 4A**). To digest the RNA on the cells, 10^7^ cells were washed once by HBSS and resuspended by 1 ml of HBSS, then added with 0, 1 or 5 μl of RNase cocktail at 37 °C for 10 mins.

### Copper-free click conjugation to RNA

Copper-free conditions were used in all experiments to avoid copper in solution during the conjugate of biotin to the azido sugars. All experiments used dibenzocyclooctyne-PEG4-biotin (DBCO-biotin, Sigma) as the alkyne half of the cycloaddition. To perform the SPAAC, RNA in pure water was mixed with 1x volumes of “dye-free” Gel Loading Buffer II (df-GLBII, 95% Formamide, 18 mM EDTA, and 0.025% SDS) and 500 μM DBCO-biotin. Typically, these reactions were 30 μL df-GLBII, 27 μL RNA, 3 μL 10 mM stock of the DBCO reagent. Samples were conjugated at 55°C for 10 min to denature the RNA and any other possible contaminants. Reactions were stopped by adding 2x volumes (120 μL) of RNA Binding Buffer (Zymo), vortexing, and finally adding 3x volumes (180 μL) of 100% EtOH and vortexing. This binding reaction was purified over the Zymo column as instructed by the manufacturer and analyzed by gel electrophoresis as described below.

### RNA gel electrophoresis, blotting, and imaging

Blotting analysis of ManNAz-labeled RNA was performed according to the procedures described by Flynn et al. with the following modifications. RNA on the column was eluted by 18 μl H_2_O and then add 18 μl of df-GLBII with EB (Thermo Fisher Scientific). To denature, RNA was incubated at 55°C for 10 min and crashed on ice for 3 min. Samples were then loaded into a 1% agarose-formaldehyde denaturing gel (Northern Max Kit, Thermo Fisher Scientific) and electrophoresed at 60 V for 60 min. Total RNA was then visualized in the gel using iBright 1500 (Invitrogen). RNA transfer to NC membrane (0.45 μm) occurred as per the Northern Max protocol for 2-16 h at 25°C. After transfer, RNA was crosslinked to the NC using UV-C light (0.18 J/cm^2^). NC membranes were then blocked with Protein Free Blocking Buffer, PBS (Li-Cor Biosciences) for 60 min at 25°C. After blocking, the blot was incubated with anti-Biotin-HRP (1:1000) for 2-16 h at 4°C. Excess anti-biotin-HRP was briefly washed from the membranes by 0.1% Tween-20 (Sigma) in 1x PBS. NC membranes were imaged on iBright 1500 (Invitrogen).

For detecting the interaction of recombinant protein-Fc chimera with glycoRNAs, native RNA extracted from unlabeled cells were subjected to gel electrophoresis and blotting as described above. Recombinant protein-Fc (1.5 μg/ml) was pre-complexed with anti-Fc-HRP (1.5 μg/ml) in 5 ml FACS buffer at 4°C for 1 h. After blocking, the NC membrane was incubated with the pre-complexed protein and second antibody solution at 4°C for 2 h. Excess anti-Fc-HRP was briefly washed from the membranes by 0.1% Tween-20 (Sigma) in 1x PBS. NC membranes were imaged on iBright 1500 (Invitrogen).

### Analysis of monocyte adhesion on ECs

To perform the adhesion assay, ECs were seeded on 6-well plates (Falcon). When ECs formed a dense layer and completely covered the dish, a final concentration of 10 ng/mL TNF-α and 1 μg/ml of LPS (Sigma) were added to the medium and the ECs were cultured for an additional 16-24 hours. THP1 cells were labeled by 1 μM PHK67 (Sigma) according to the manufacturer’s instruction. 10^7^ THP1 cells or THP1-RFP cells were spin down and washed once by HBSS with Ca^2+^ and Mg^2+^ (Gibco). THP1 or THP1-RFP cells were then incubated with 1 mL HBSS with Ca^2+^ and Mg^2+^ supplemented with 0, 1 or 5 μl of RNase cocktail for at 37°C for 10 minutes. THP1 cells were labeled by 1 μM PHK67 (Sigma) according to the manufacturer’s instruction. 10^6^ Labeled THP1 cells or THP-1-RFP cells were added into each well and incubated for 1 h. Unattached or loosely attached THP1 cells were washed twice by 2 ml HBSS with Ca^2+^ and Mg^2+^. Attached THP1 cells were fixed by adding 4% formaldehyde for 15 mins at room temperature. The cells then were imaged by microscope (Evos FL Auto 2, Invitrogen). At least 6 images were taken from each well and adhesion cells were counted by Qupath.

### Fluorescence-activated cell sorting (FACS) and microscopy analysis

THP1 cells were grown as described above and spin down by centrifuge. Cells were resuspended in FACS Buffer (0.5% bovine serum albumen (Thermo Fisher Scientific) in 1x PBS), counted, and aliquoted to 1,000,000 cells per 200 μL FACS Buffer, incubating on ice for 30 min to blocking. For RNase digestions, 1 μl of RNase cocktail was added to the blocking buffer and incubated at 37°C for 10 min and then put on ice for 20 min. After blocking, cells were brought to 25°C for 5 min, then spun for 5 min at 4°C and 350x g. Cells were washed once with 150 μL FACS Buffer and spun as above.

1.5 μg/ml recombinant protein-Fc chimera was precomplexed with 1.5 μg/ml anti-human Fc-FITC in 200 μl FACS buffer on ice at dark for 1 h. Then the THP1 cells were resuspended with 200 μl of the pre-complexed recombinant protein-Fc-Secondary solution and incubated on ice in the dark for 30 min, washed once with 200 μl FACS buffer and resuspended in 200 μl FACS buffer. 150 μl of cells were sent to our FACS core for analysis and the other 50 μl of cells was fixed by adding 50 μl of 4% formaldehyde for 15 min at room temperature. The fixed cells were spread onto 12 well plate and centrifuged for 5 min. Then the cells were imaged by microscopy (Evos FL Auto 2, Invitrogen).

### Immunoprecipitation

50 μL anti-Fc beads were pro-conjugated with 10 μg of recombinant protein-Fc chimera in samples buffer (50 mM Tris, pH 7.4, 150 mM NaCI, 0.5% Triton X-100) for 2 hours at 4°C. After washing three times with samples buffer, beads were incubated with 10 μg of biotinylated RNA from ManNAz-labeled THP1 cells for 2 hours at 4°C. After washing three times with samples buffer, the beads were suspended in 50 μL RNA binding buffer and heated for 5 minutes at 50°C. Remove the beads and transfer the RNA extract solution to a new tube. Here, 100 μL of pure water was added and vortexed for 10 seconds, and then 300 μL of 100% ethanol was added and vortexed for 10 seconds. The RNAs were purified over a Zymo column followed by northern blot analysis as described above.

### Purification of glycoRNA-L and glycoRNA-S

The purification of biotin-labeled glycoRNAs was achieved by streptavidin beads with the following steps: 20 μL of MyOne C1 Streptavidin beads (Thermo Fisher Scientific) per reaction were blocked with 50 ng/μL glycogen (Thermo Fisher Scientific) and 1 U/μL RNase Inhibitor (NEB) in Biotin Wash Buffer (10 mM Tris HCl pH 7.4, 1 mM EDTA, 100 mM NaCl, 0.05% Tween-20) for 1 hour at 25°C. The beads were washed by Biotin Wash Buffer and the spun down. Next, 50 μg of the biotinylated total RNAs were diluted in 1 mL Biotin Wash Buffer and incubated with the blocked MyOne C1 beads for 4 hours at 4°C. Beads were washed to remove un-bound RNAs for two times with 1 mL of ChIRP Wash Buffer (2× SSC, 0.5% SDS), then washed two time with 1 mL of Biotin Wash Buffer each, followed by two time washes with 1 mL NT2 Buffer (50 mM Tris HCl, pH 7.5, 150 mM NaCl, 1 mM MgCl2, 0.005% NP-40). All washes were performed at room temperature for 3 minutes each. The streptavidin beads were then eluted by incubation in 1 mL TRIzol at 25°C for 10 minutes and the glycoRNA was extracted and purified by Zymo C column. To separate glycoRNA-L and glycoRNA-S, first the purified glycoRNAs were loaded on low-temperature melting agarose gel and separated by electrophoresis. The gel slices containing glycoRNA-L and glycoRNA-S were collected. 4 volumes of RNA binding buffer was added into the tube containing gel slices and incubated at 50°C for 10 min. Then added 1 volume of 100% ethanol and incubated at 50°C for additional 5 min. The glycoRNA-L and glycoRNA-S were purified by Zymo column as described above.

### Western blot

Western blot was performed as described previously (22). Briefly, Cells were quickly rinsed with ice-cold PBS, and directly lysed with RIPA buffer (150 mM NaCl, containing proteinase inhibitor cocktail 1 and 3 and phosphatase inhibitor cocktail (Cell Signaling Technology) on ice for 15 minutes. After centrifugation at 12,000x g for 15 minutes at 4°C, lysates were heated at 95°C for 10 minutes in 1x NuPAGE LDS loading buffer (ThermoFisher Scientific) containing 5 mM DTT. Samples were then resolved by SDS-PAGE using AnyKD Criterion TGX Precast Midi Protein Gels (Bio-Rad Laboratories) and transferred to nitrocellulose membranes. Membranes were blocked in blocking buffer, and incubated with primary antibodies (diluted in blocking buffer) at 4°C overnight. After washing three times for 3 minutes each in 1x PBS with 0.1% Tween-20 (PBST), membranes were incubated with secondary antibodies at room temperature for 60 minutes, followed by the same 3x PBST washing. Membranes were finally rinsed in 1x PBS and scanned on iBright 1500 (Invitrogen). Primary antibodies used: mouse monoclonal anti-hSiglec-5 (R&D systems, 1:1000), rabbit anti-VCAM-1 polyclonal antibody (Santa Cruz, 1:1000), rabbit polyclonal anti-β-actin (Santa Cruz, 1:1000). Secondary antibodies used: goat anti-rabbit IgG-HRP (Invitrogen, 1:1000) and rabbit anti-mouse IgG-HRP (Invitrogen, 1:1000).

### Cell viability

The cell viability was tested by Trypan Blue Exclusion Test. Briefly, 10^7^ THP1 cells were spun down and then resuspended into 1 ml HBSS buffer and incubated with 0, 1 or 5 μL of RNase cocktail at 37°C for 10 min. The cells then were mixed with equal volume of 0.4% Trypan blue (gibco) and counted by a cellometer (Nexcelom Bioscience). The cell viability was presented as all cell numbers-stained cell numbers/all cell numbersX100%.

### Statistics

Student’s t-test (unpaired, unequal variance) was used to assess experimental significance, unless specified otherwise.

## SUPPLEMENTAL INFORMATION

Supplemental information is included.

## ACKNOWLEDGMENTS

We thank Dr. Daping Fan for providing THP1 and THP1-RFP cells. We thank Dr. Kun Cheng for providing HeLa cells and Dr. Shizhen Wang for providing HEK293 cells. M.F. is partly supported by UMKC Funding for Excellence grant and SPiRE.

## AUTHOR CONTRIBUTIONS

M.F. conceived, designed and supervised the overall study. M.F. performed most of the molecular experiments. Y.Q performed the tissue expression pattern of glycoRNAs in mice. E.H. and Z.S. performed part of the monocyte adhesion studies. T.L performed western blot analysis and lab management. H.T and K.T assisted in the experiments, and M.F. wrote the manuscript.

## DECLARATION OF INTERESTS

The authors declare no competing interests.

## Supplemental Information

**Figure S1.**
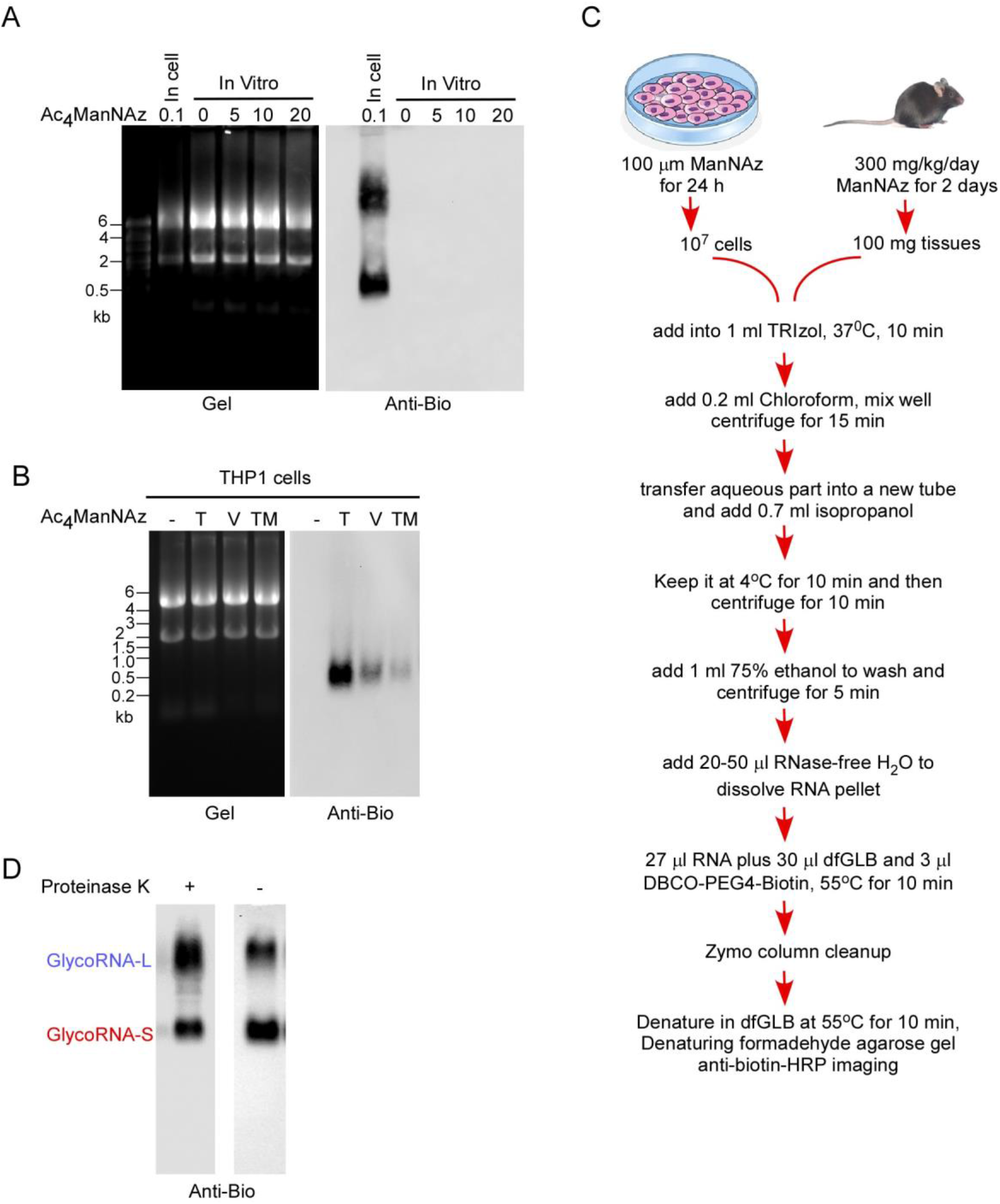
Identification of glycoRNA-L and glycoRNA-S in THP1 cells, related to Figure 1. (A) Blotting of in cellular or in vitro Ac_4_ManNAz-labeled RNA. Cells were treated with 100 μM Ac_4_ManNAz for 24 hours while native RNA in vitro was treated with 0, 5, 10 and 20 mM of Ac4ManNAz at 37 °C for 2 hours. (B) THP1 cells were treated with Ac_4_ManNAz from Tocris (T), Vector Labs (V) and TargetMol (TM). RNAs were extracted from the cells and analyzed by northern blot. Representative images are shown. (C) Schematic of RNA extraction protocol. Total RNAs were always extracted by TRIzol and precipitated by isopropanol. (D) Blotting of RNA treated with or without proteinase K and then purified by Zymo column.

**Figure S2.**
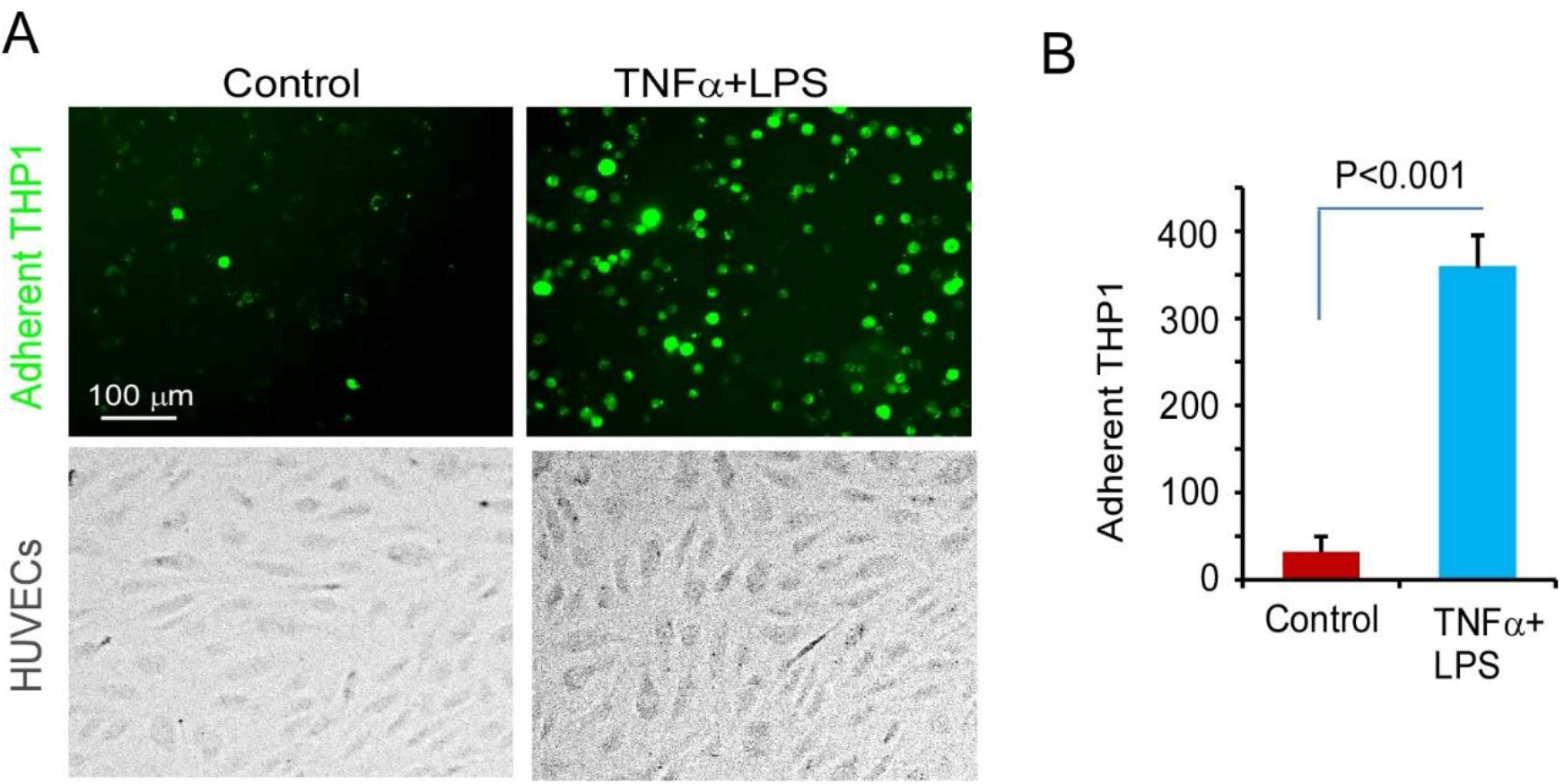
THP1 cells adherent to activated HUVECs but not rest HUVECs, related to Figure 2. (A) The cell images taking fromTHP1 cells adherent to activated HUVECs. HUVECs were treated with or without 10 ng/ml TNFα and 1 μg/ml LPS for 24 hours. THP1 cells were labeled by PHK67 for 5 min and then incubated with HUVECs at 37 °C for 1 hour. The images were taken from 6 different fields. Representative image represents one fourth of original image to make the cells more visible. (B) 6 fields per well were imaged and the cell numbers in each field were counted by Qupath. The adherent cell numbers were averaged and presented as mean±SD, n=3.

**Figure S3.**
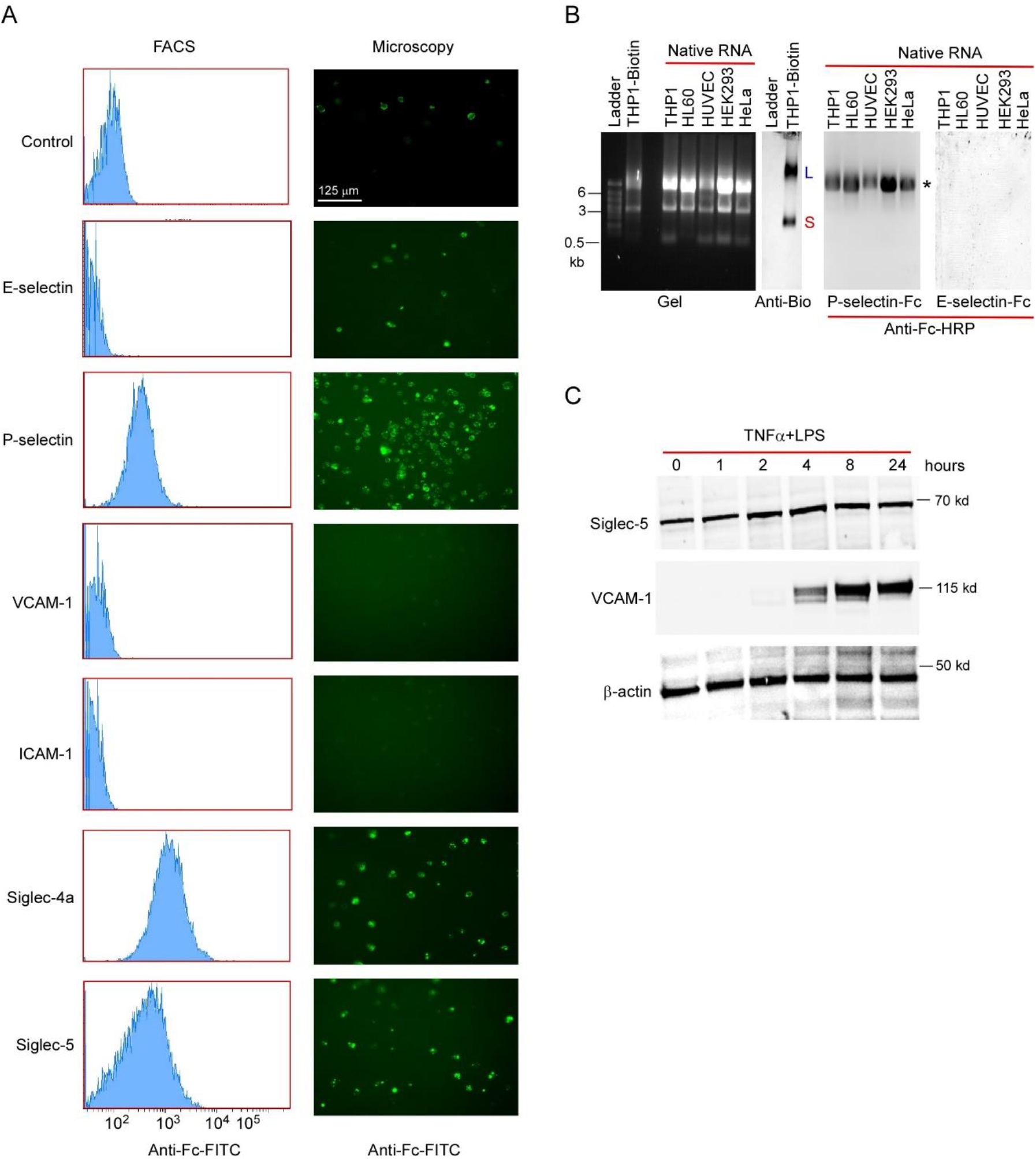
Searching the binding partner of glycoRNA-L and glycoRNA-S, related to Figure 3. (A) FACS analysis of THP1 cells labeled by pre-complex of E-selectin-Fc, P-selectin-Fc, VCAM-1-Fc, ICAM-1-Fc, Siglec-4a-Fc, Siglec-5-Fc and anti-Fc-FITC (left). The images taken from these cells by microscopy (right). (B) Total native RNAs were extracted from the cell lines as indicated. 20 μg RNA from each cell line were blot on NC membrane as described above. After blocking, the membrane was probed by pre-complex of P-selectin-Fc (1.5 μg/ml) or E-selectin-Fc (1.5 μg/ml) with anti-Fc-HRP (1.5 μg/ml) for 2 h at 4°C. The blot was imaged by iBright 1500. *non-specific binding. Blot of biotinylated RNA from labeled THP1 cells was probed by anti-biotin-HRP. (C) Western blot of cell lysates from HUVECs treated with TNFα (10 ng/ml) and LPS (1 μg/ml) for different time points as indicated. The blot was probed by primary antibodies as indicated and imaged by secondary antibody-HRP.

## Notes

### Competing Interest Statement

The authors have declared no competing interest.

